# Histones and nucleoid-associated proteins in *Methanosarcina acetivorans* mediate recovery from stress and stasis

**DOI:** 10.1101/2025.11.24.690127

**Authors:** Alienor S. Baskevitch, Dipti D. Nayak

## Abstract

Many archaea encode both histones and **<u>N</u>**ucleoid-**<u>A</u>**ssociated **P**roteins (NAPs), which we refer to collectively as archaeal **<u>D</u>**NA **<u>B</u>**inding **<u>P</u>**roteins (DBPs). Whether these DBPs jointly work to compact the genome or have distinct functions remains unknown. Here, we have developed the methanogen, *Methanosarcina acetivorans,* as a platform to study the function of archaeal DBPs *in vivo. M. acetivorans* encodes one archaeal histone, *hmaA,* and two copies of the archaea-specific NAP, *mc1* (*mc1a* & *mc1b).* We found that each DBP is individually dispensable, but at least one copy of *mc1* appears to be required for growth. The growth of the single and double DBP deletion mutants was, by and large, like the parent strain under optimal growth conditions. However, after exposure to stresses or extended periods of incubation in stationary phase, the DBP deletion strains often recovered growth much faster than the parent strain. Conversely, over-expression of DBPs led to a delay in growth recovery that could be abrogated by introducing point mutations in DNA-binding residues. Together, our data suggest that histone and archaea-specific NAPs have partially overlapping roles in *M. acetivorans* and likely protect the genome after exposure to stress or during prolonged periods of growth stasis. Our findings emphasize that there is no unified function for histones across the tree of life and instead imply that archaeal histones join the ranks of other archaeal NAPs in having strain-specific functions.

**Importance:** Though it is known that many archaea encode histones in tandem with archaea-specific **<u>N</u>**ucleoid-**<u>A</u>**ssociated **<u>P</u>**roteins (NAPs), the interplay between these two classes of **<u>D</u>**NA-**<u>B</u>**inding **<u>P</u>**roteins (DBPs) *in vivo* is not known. Most studies on archaeal DBPs have focused either on archaea known to form histone-based chromatin, or on strains which lack histones and compact their genomes exclusively with archaea-specific NAPs. Our study, therefore, fills an important gap in the literature by characterizing DBPs in a model methanogen, *Methanosarcina acetivorans*, which encodes both histones and archaea-specific NAPs. While no DBP is essential, at least one copy of MC1 appears to be needed, likely for genome compaction. In addition, mutants lacking DBPs had a growth advantage after being subject to stresses or long periods of growth stasis, indicating that DBPs likely function in genome maintenance when growth has stalled or stopped.

## Introduction

Proteins that shape chromosome architecture vary across all domains of life^1^. Bacteria use small, basic proteins to induce topological changes - such as bending, bridging, and wrapping - that ultimately lead to chromosome compaction^2,3^. These proteins are called **<u>N</u>**ucleoid-**<u>A</u>**ssociated **<u>P</u>**roteins (NAPs) as they are primarily localized to the bacterial nucleoid. In contrast, eukaryotes use histones to organize genomic DNA into chromatin. While NAPs and histones are primarily associated with genome compaction, it is now well-established that they also play other important cellular functions. Bacterial NAPs are involved in DNA protection, chromosome segregation, stress response and gene regulation^2,4^. Likewise, histones play a role in transcriptional regulation^5^ and have recently been shown to also function as a copper reductase enzyme in eukaryotes^6^. Archaea often encode histones^7^ as well as NAPs^8^ and the cellular roles of these two proteins are not well established, especially when they co-occur. Archaeal histones are structurally and functionally related to their eukaryotic counterparts^5,7,9–11^. In some cases, archaeal histones have been demonstrated to form nucleosomes *in vitro*^12,13^ and *in vivo*^14,15^. Indeed, histones are essential in thermophilic archaea, like *Thermococcus kodakerensis,* which is consistent with their role in genome compaction *in vivo*. However, histones are not universally conserved across archaea^8^. Also, histones are dispensable in halophilic archaea like *Halobacterium salinarum* and *Haloferax volcanii*, where they have been proposed to function more like transcription factors^16,17^. In *Methanosarcina* spp., histones are also not essential and have been proposed to play an auxiliary role during DNA-damage repair^18–22^. In cases where histones are absent or non-essential, archaeal NAPs, such as Cren7 and Alba, have been implicated in genome compaction instead^1,18,23–25^. In archaea that encode histones and NAPs, the functional demarcation between these two groups of proteins is not well defined.

Here, we use the genetically tractable mesophilic methanogen, *Methanosarcina acetivorans*, which encodes one copy of an archaeal histone and two copies of the archaea-specific NAP, **<u>m</u>**ethanogen **<u>c</u>**hromosomal protein <u>1</u> (MC1), as a model system to delineate the role of histones and NAPs *in vivo.* We refer to histones and NAPs collectively as DNA-binding proteins (DBPs) hence forth, as this is the only feature that unifies these otherwise distinct groups of proteins. To analyze the function of each DBP, we attempted to generate mutants lacking DBPs in all possible combinations. All three DBPs in *M. acetivorans* are individually dispensable. However, we were unable to recover a mutant lacking both copies of MC1, suggesting that at least one copy of this NAP is essential, perhaps for genome compaction. Under standard growth conditions, a few DBP deletion mutants had a minor growth defect. Remarkably, the absence of DBPs speeds up the rate of recovery after cells are subject to stressors or prolonged periods of growth stasis. Our observations are consistent with a protective role for DBPs under stress or during periods of quiescence, which is likely how these cells spend majority of their time in the natural environment.

## Results

### All three DNA-binding Proteins are dispensable in *Methanosarcina acetivorans*

All three DBPs in *M. acetivorans* are encoded as monocistronic genes (Fig. S1). Sequence-and structure-alignment suggest that these DBPs have maintained all the critical residues implicated in binding DNA (Fig. S1). A previous study showed that the histone from *Methanosarcina mazei* Gö1 (HMm), while dispensable, is important for optimal growth^18^. To the best of our knowledge, mutational studies of the *mc1* gene(s) have not been reported. Hence, we chose to delete each DBP in *M. acetivorans* individually, and in all possible combinations, using our well-established CRISPR-Cas9 genome-editing methodology^26^. We found that all three DBPs in *M. acetivorans* are individually dispensable (Fig. 1). Whole genome sequencing revealed that none of these single knockout mutants had any other off-target or suppressor mutations (Table S1). During growth on a single substrate, either methanol, trimethylamine (TMA), or acetate, some of the single knockout strains had a mild growth disadvantage (Table 1). For instance, the Δ*hmaA* mutant grew 20% slower than the parent strain (WWM60; referred to as wildtype or WT henceforth) on acetate and all three mutants had a 20% longer lag when switching from methanol to TMA (Fig. S3; Table 1). To ensure that the growth phenotypes did not stem from changes in cell shape or size, we measured these parameters for all single knockouts and did not observe any significant changes relative to the parent strain (Fig. S4). We were readily able to obtain Δ*hmaAΔmc1a* and Δ*hmaAΔmc1b* double mutants and their growth phenocopied the WT strain (Figs. 1 and S5; Table 1). Intriguingly, we were unable to obtain a double mutant that lacks both MC1a and MC1b or the triple knockout mutant, even after several attempts (as outlined in the Methods and Table S2). These results suggest that, while no individual DBP is essential, at least one copy of *mc1* appears to be essential in *M. acetivorans*.

**Figure 1:**
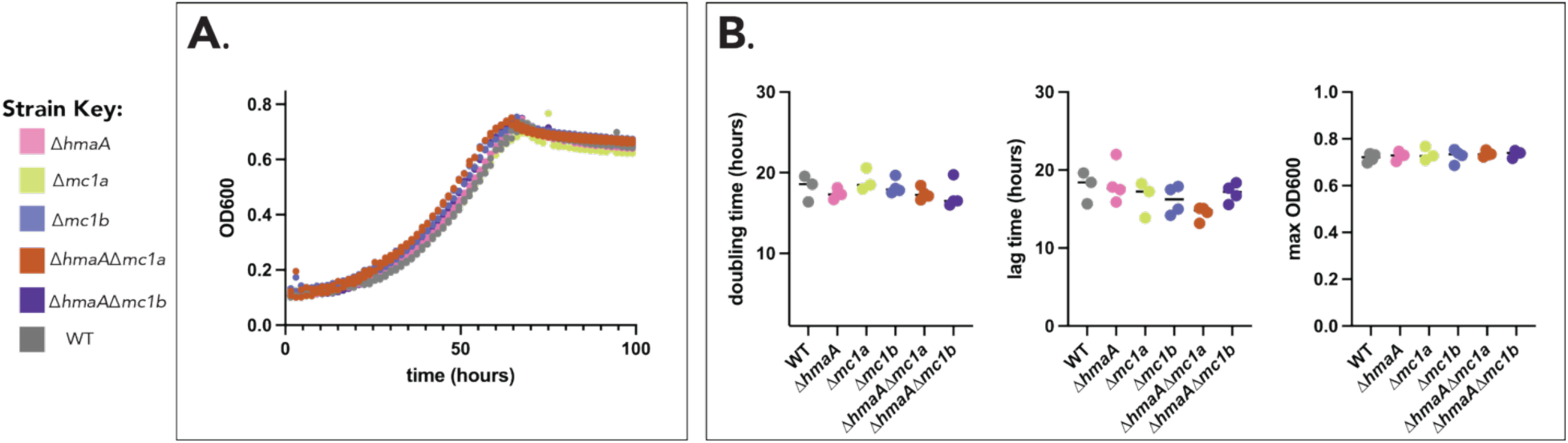
DNA-binding proteins (DBPs) are not individually essential in *M. acetivorans.* **A>** Growth curves of DBP deletion mutants (colored circles) and the parent strain (WWM 60; denoted as wildtype or WT; gray circles) measured as a change in optical density (at 600 nm; OD600) over time in high-salt (HS) minimal medium supplemented with 50 mM trimethylamine as the growth substrate. Representative growth curves for at least two replicates are shown. **B.** Quantification of doubling time (in hours), lag time (in hours), cell yield (measured as maximum OD600) for at least three replicates of each strain shown in panel **A**. The horizontal line represents the mean of the replicates. No significant differences from WT in doubling time, cell yield, and lag time were observed using a one-way ANOVA with Dunnett’s multiple comparisons test. See Table 1 for quantification of growth parameters from these growth curves.

**Table 1:** Growth data for DBP deletion strains under standard laboratory growth conditions. Doubling time, lag time, and final optical density at 600 nm (representative of growth yield) are reported for each strain under the conditions tested. All values are reported as the mean ± one standard deviation (SD). Data represent the mean of at least three replicates. For values that are significantly different from the parent strain (WWM60 denoted as wildtype or WT) for a given condition (one-way ANOVA, Dunnett’s multiple comparisons test), percentage change from the WT is reported; sign (+ or -) indicates whether the mutant showed an increase or reduction relative to WT, respectively. Substrates marked with an asterisk (*) denote growth experiments that were conducted in a 24-well plate with an automated plate reader (see Methods). All other growth curves were conducted in anaerobic Balch tubes.

| strain | substrate | doubling time (hours) | significantly different than WT (p<0.05)? | lag time (hours) | significantly different than WT (p<0.05)? | final OD (OD600) | significantly different than WT (p<0.05)? |
| --- | --- | --- | --- | --- | --- | --- | --- |
| WT | TMA* | 18.2±1.6 | n/a | 17.9±2.0 | n/a | 0.72±0.02 | n/a |
| $\Delta hmaA$ | | 17.3±0.6 | no | 18.3±2.6 | no | 0.73±0.02 | no |
| $\Delta mc1a$ | | 19.0±1.4 | no | 16.4±2.3 | no | 0.73±0.03 | no |
| $\Delta mc1b$ | | 18.3±1.0 | no | 16.1±1.8 | no | 0.73±0.03 | no |
| $\Delta hmaA\Delta mc1a$ | | 17.4±0.8 | no | 14.5±0.9 | no | 0.74±0.01 | no |
| $\Delta hmaA\Delta mc1b$ | | 17.2±1.7 | no | 17.1±1.2 | no | 0.74±0.01 | no |
| WT | MeOH | 10.0±0.2 | n/a | 6.6±0.8 | n/a | 1.09±0.02 | n/a |
| $\Delta hmaA$ | | 10.4±0.2 | no | 6.2±0.3 | no | 1.11±0.02 | no |
| $\Delta mc1a$ | | 10.6±0.2 | no | 5.8±0.6 | no | 1.09±0.02 | no |
| $\Delta mc1b$ | | 10.0±0.1 | no | 6.4±0.4 | no | 1.10±0.03 | no |
| WT | acetate | 47.4±3.0 | n/a | 127.3±21.2 | n/a | 0.21±0.01 | n/a |
| $\Delta hmaA$ | | 56.7±3.2 | yes, +19.6% | 144.8±24.2 | no | 0.20±0.01 | no |
| $\Delta mc1a$ | | 47.5±2.6 | no | 131.0±18.5 | no | 0.21±0.00 | no |
| $\Delta mc1b$ | | 49.6±1.7 | no | 105.5±19.0 | no | 0.212±0.01 | no |
| WT | TMA, after passaging on MeOH | 13.7±0.7 | n/a | 86.5±4.7 | n/a | 1.23±0.01 | n/a |
| $\Delta hmaA$ | | 12.3±0.5 | yes, -10.0% | 103.0±2.2 | yes, +24.2% | 1.25±0.01 | no |
| $\Delta mc1a$ | | 13.3±0.6 | no | 100.6±6.8 | yes, +18.5% | 1.26±0.01 | yes, +2.7% |
| $\Delta mc1b$ | | 11.1±0.1 | yes, -19.9% | 103.5±1.7 | yes, +18.9% | 1.21±0.02 | no |
| WT | acetate | 49.1±3.4 | n/a | 80.1±14.9 | n/a | 0.16±0.01 | n/a |
| $\Delta hmaA\Delta mc1a$ | | 53.8±2.0 | no | 76.9±4.3 | no | 0.16±0.01 | no |
| $\Delta hmaA\Delta mc1b$ | | 55.5±4.6 | no | 78.8±1.5 | no | 0.16±0.02 | no |

### Each DBP in *M. acetivorans* has a unique transcriptomic profile

To test if the dispensability of individual DBPs stems from functional redundancy, we obtained the global transcriptomic profile of the three single knockout mutants and WT in minimal medium supplemented with 50 mM TMA. We chose growth on TMA for this experiment because the growth parameters of all single knockout strains were nearly identical under these conditions (Table 1), which would minimize any impact of altered growth parameters on the transcriptome. All three DBPs were expressed in WT grown on TMA but their expression levels varied substantially. The *mc1a* locus was expressed 3-fold more than *mc1b*, and *hmaA* was the least expressed DBP, *ca.* 3-fold lower than *mc1b* (Fig. S6). In the single knockout background, none of the remaining DBPs changed in expression (Fig. S6), which precluded the likelihood of functional compensation at the transcriptional level.

Next, we explored how each DBP impacts the global transcriptional profile of the cell by comparing the transcriptome of the corresponding deletion strain to WT. We designated a gene as being differentially expressed if it had a log_2_ fold change of greater than 0.1 and was statistically significant (q-value < 0.05). At a global level, each DBP has a unique transcriptional profile (Fig. 2A, Table S3). Nearly 10% of the genes in the genome (405) were differentially expressed (DE) in the *Δmc1b* knockout mutant. This was in sharp contrast to the *Δmc1a* strain, where only 12 genes were DE. A response regulator (MA_RS14075) was the only protein-coding gene differentially expressed in both the Δ*mc1a* and *Δmc1b* mutant (Table S3). Thus, even though our genetic analyses suggest that the two *mc1* genes have some functional overlap (Table S2), it is not evident in the global transcriptome. 61 genes were DE in the *ΔhmaA* strain (Fig. 2A, Table S3), and 30% of these were shared with at least one of the Δ*mc1* deletion mutants. Across all mutants, the DE genes did not localize to any region of the genome or to either the + or – strand (Fig. 2B). The DE genes were not enriched for any specific functional category (Fig. S7 and Tables S4- S8). A TTTTC motif was enriched in the promoter region of DE genes for the Δ*mc1b* mutant, which is consistent with previous findings that this protein preferentially binds certain DNA sequences^27,28^ (Fig. S8). In contrast, no such motifs were found for DE genes in the Δ*hmaA* and Δ*mc1a* mutant (Fig. S8).

**Figure 2:**
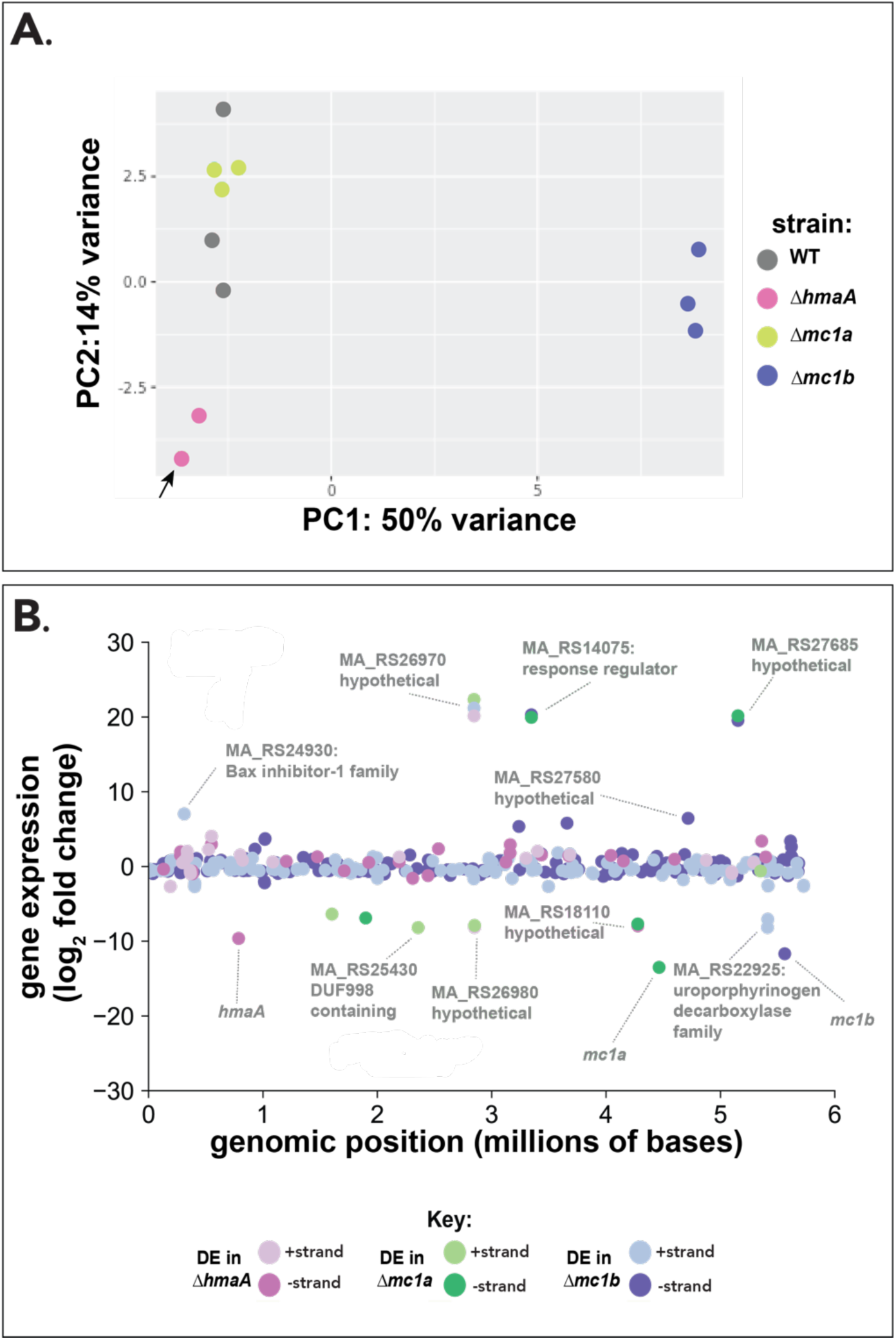
DNA-binding protein (DBP) deletion mutants have a unique and distinct transcriptional profile compared to the parent strain. **A.** Principal Component Analysis (PCA) plot showing transcriptional profiles of the three DBP single deletion mutants (colored circles) and the parent strain (WWM 60; denoted as wildtype or WT; gray circles) grown in high-salt (HS) minimal medium with 50 mM trimethylamine (TMA). Black arrow indicates two points that overlap. **B.** Scatter plot depicting genomic location and strand information for differentially expressed (DE) genes in the single DBP deletion strains. Pink, green, and blue circles denote genes that are differentially expressed in *ΔhmaA*, *Δmc1a, Δmc1b* mutants. Strand information is given by the shade of the circle: light circles show genes on the positive strand, while dark circles show genes on the negative strand. Genes with a large fold-change (>log_2_ 7.5-fold) in any of the mutants are labeled with locus tags and annotation information. Genes that are DE in multiple mutants are labeled with the colors of each mutant strain in which the gene is DE. DE tRNAs are not depicted in this figure but are included in **Table S3**.

### Some DBPs impact growth after exposure to stressors

Unlike the natural environment, laboratory conditions are optimized for fast growth and are devoid of abiotic or biotic stressors that DBPs might specifically respond to. Indeed, the histone deletion mutant of *M. mazei* Gö1 was shown to be more sensitive to UV shock than the parent strain^18^. To test the role of DBPs in stress response, we measured the growth phenotype of the single and double DBP deletion mutants in the presence of a known DNA-damaging chemical agent (mitomycin C; MMC), at high temperature (40°C), or after UV exposure. In all cases, the DBP knockout mutants were viable (Figs. 3 and S9, Table 2). After exposure to 1μg/mL MMC, a few mutants had a mild but significant growth defect. For example, the *Δmc1b* mutant and the Δ*hmaAΔmc1b* mutant had a 20% and 40% longer lag phase, respectively (Fig. 3A, Table 2). At elevated temperatures, the *ΔhmaAΔmc1a* mutant grew 37% slower than WT (Fig. 3B, Table 2). However, sometimes strains lacking *hmaA* and/or *mc1b* also had a growth advantage. For example, at 40°C, the lag time for the *Δmc1b* mutant was 39% shorter than WT (Fig. 3B). The *ΔhmaA* and *Δmc1b* single knockout mutants also recovered 18% and 26% faster than WT after exposure to UV shock, respectively (Fig. 3C). This growth advantage was further amplified in the *ΔhmaAΔmc1b* double mutant, which recovered 35% faster than WT after UV exposure (Fig. 3C).

**Figure 3:**
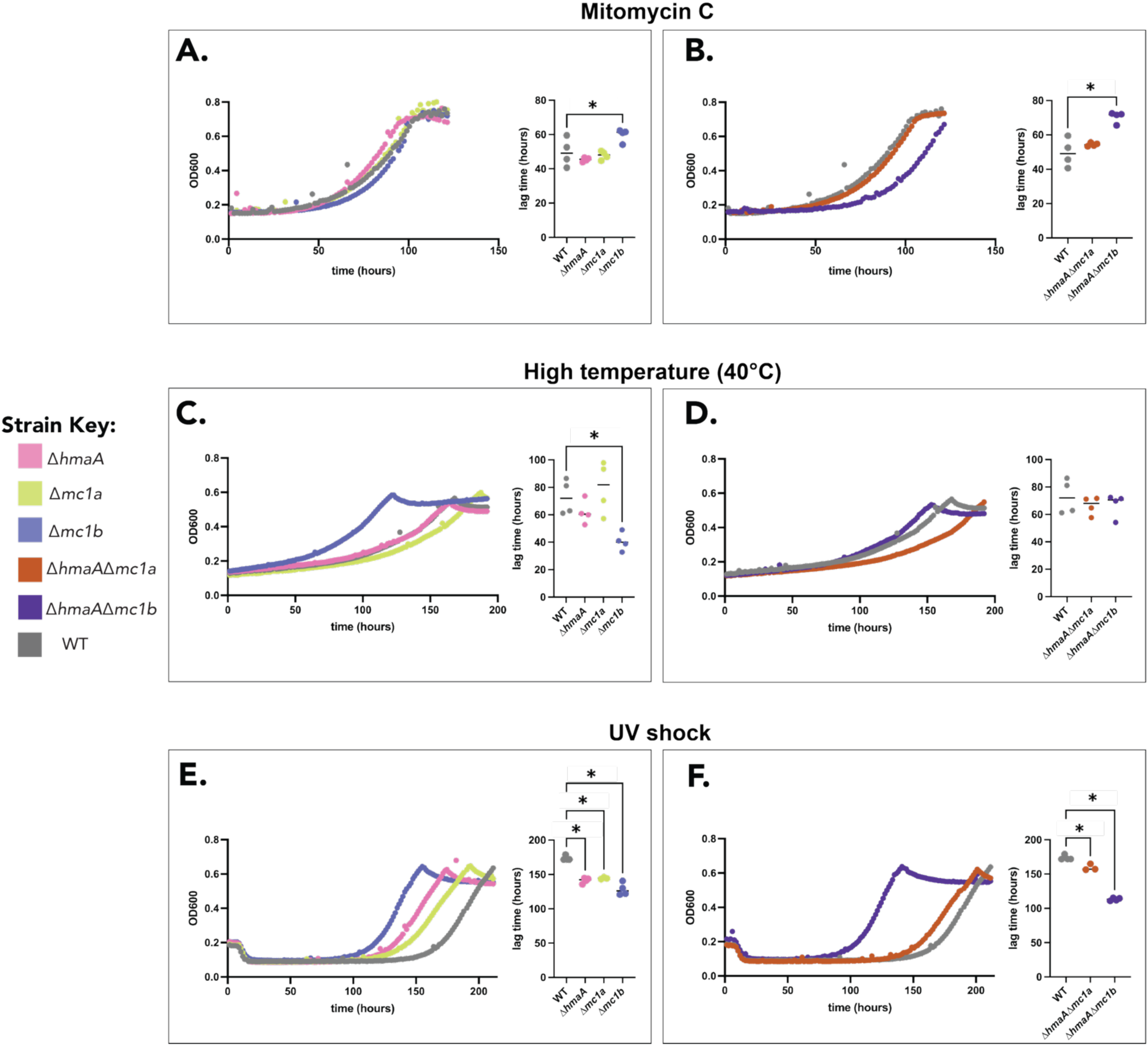
Growth of DNA-binding proteins (DBP) deletion mutants after the application of an abiotic stress. Growth curves of DBP deletion mutants (colored circles) and the parent strain (WWM 60; denoted as wildtype or WT; gray circles): **A.** and **B.** in media supplemented with 1 μM Mitomycin C (MMC); **C.** and **D.** at elevated temperature (40°C); and **E.** and **F.** following UV shock (see Methods) and transfer to recovery media. Each condition is split to depict single DBP deletion strains on the left (panels **A, C,** and **E)** and double DBP deletion strains on the right (panels **B, D,** and **F)**. Growth is measured as a change in optical density (at 600 nm; OD600) over time in high-salt (HS) minimal medium supplemented with 50 mM trimethylamine as the growth substrate. A representative growth curve for each strain is shown. The lag time (in hours) was quantified for least three replicates of each strain shown. The horizontal line represents the mean of the replicates. Statistically significant differences relative to WWM60 (p<0.05) based on a one-way ANOVA with Dunnett’s multiple comparisons test are depicted as follows: * denotes p-value ≤ 0.05. See Table 2 for quantification of the growth parameters shown in this figure.

**Table 2:** Growth data for DBP deletion strains in the presence of stressors. Doubling time and lag time are reported for each strain under the conditions tested. For UV shock, growth following a one-time exposure to UV (see Methods for details) was measured. For high temperature and Mitomycin C, cultures were exposed to the stress for the duration of the growth experiment (see Methods for details). Data represent at least three replicates. Mean values and standard deviations for each growth parameter are denoted. For values that are significantly different from the WT strain for a given condition (one-way ANOVA, Dunnett’s multiple comparisons test), percentage change from the WT is reported; sign (+ or -) indicates whether the mutant showed an increase or reduction relative to WT, respectively. All growth curves were conducted in a 24-well plate with an automated plate reader (see Methods) using high-salt (HS) minimal medium supplemented with 50mM TMA.

| strain | stress applied | doubling time (hours) | significantly different than WT (p<0.05)? | lag time (hours) | significantly different than WT (p<0.05)? |
| --- | --- | --- | --- | --- | --- |
| WT | Mitomycin C | 29.6±2.3 | n/a | 49.6±8.2 | n/a |
| $\Delta hmaA$ | | 23.4±0.8 | yes, -21.0% | 45.4±1.2 | no |
| $\Delta mc1a$ | | 26.7±1.8 | no | 47.8±2.5 | no |
| $\Delta mc1b$ | | 23.7±4.1 | yes, -19.8% | 59.7±3.8 | yes, +20.4% |
| $\Delta hmaA\Delta mc1a$ | | 25.7±0.5 | no | 54.4±1.0 | no |
| $\Delta hmaA\Delta mc1b$ | | 27.8±3.1 | no | 70.5±3.3 | yes, +42.0% |
| WT | high temperature stress (40°C) | 48.5±6.9 | n/a | 72.9±12.7 | n/a |
| $\Delta hmaA$ | | 60.8±11.7 | no | 61.8±8.7 | no |
| $\Delta mc1a$ | | 54.3±11.9 | no | 79.7±19.2 | no |
| $\Delta mc1b$ | | 41.8±3.2 | no | 40.4±6.7 | yes, -39.4% |
| $\Delta hmaA\Delta mc1a$ | | 66.2±4.6 | yes, +36.6% | 66.3±6.6 | no |
| $\Delta hmaA\Delta mc1b$ | | 40.3±1.4 | no | 67.0±8.6 | no |
| WT | UV stress | 18.8±0.5 | n/a | 173.6±3.6 | n/a |
| $\Delta hmaA$ | | 19.2±1.9 | no | 141.3±4.1 | yes, -18.6% |
| $\Delta mc1a$ | | 21.3±0.3 | no | 144.4±1.4 | yes, -16.8% |
| $\Delta mc1b$ | | 18.1±1.0 | no | 128.4±9.1 | yes, -26.1% |
| $\Delta hmaA\Delta mc1a$ | | 19.9±1.2 | no | 159.5±4.7 | yes, -8.1% |
| $\Delta hmaA\Delta mc1b$ | | 17.9±1.3 | no | 113.12±2.1 | yes, -34.8% |

### DBPs modulate timing of exit from stationary phase

A notable phenotype from our stress-response assays was that mutants lacking HmaA and/or MC1b could recover faster after exposure to certain stressors (Fig. 3). We postulated that this phenotype could either stem from the activation of a stress-response mechanism or an override of growth arrest caused by the DBPs. To decouple these two effects, we measured the growth of DBP deletion strains after extended incubation at stationary phase i.e., after cells had ceased growing in batch culture, without any application of abiotic stress (Fig. S10). After experiencing stationary phase for *ca.* 24 hours, all strains behaved identically (Fig. 1). However, after prolonged incubation at stationary phase i.e., after 21 days, most of the DBP mutants recovered substantially faster than the WT (Fig. 4 and S11, Table 3). To control for cell viability, we measured factor 420 (F_420_) fluorescence. F_420_ is a redox cofactor that emits light in the blue-green range and is present in the cytosol of cells that have maintained intact membranes^29,30^. We observed no difference in the number of intact cells between WT and the DBP mutants after 21 days of incubation at stationary phase using F_420_ fluorescence as a readout (Fig. S12, Table S9). This outcome suggests that the DBPs may be directly responsible for modulating growth recovery from stationary phase. To validate this hypothesis, we overexpressed each DBP on a plasmid with an autonomous origin of replication using the P*mcrB*(tetO4) inducible promoter^31^, in its respective single deletion background. We confirmed expression *in trans* by immunoblotting for a FLAG tag at the C-terminus of the protein (Fig. S13). When DBPs are overexpressed, we observed a statistically significant delay in exit from stationary phase after 21 days of incubation (Figs. 4 and S14, Table 3). To test if the role of DBPs in growth recovery from stationary phase is contingent on DNA-binding, we expressed mutant alleles of *hmaA* and *mc1b* that lack residues critical for DNA binding *in vitro*^32,33^. These point mutants with diminished DNA-binding affinities were FLAG-tagged at the C-terminus and expressed in the respective knockout backgrounds (Fig. 5, Methods). By immunoblotting for the FLAG-tag, we observed that the mutant allele of each DBP was produced at levels comparable to the wildtype allele. However, the phenotype of the strains expressing mutant DBPs phenocopied the deletion mutants (Figs 5, S14; Table 4). Taken together, these data suggest that DBPs in *M. acetivorans* might bind DNA, likely to protect it when conditions are sub-optimal for growth.

**Figure 4:**
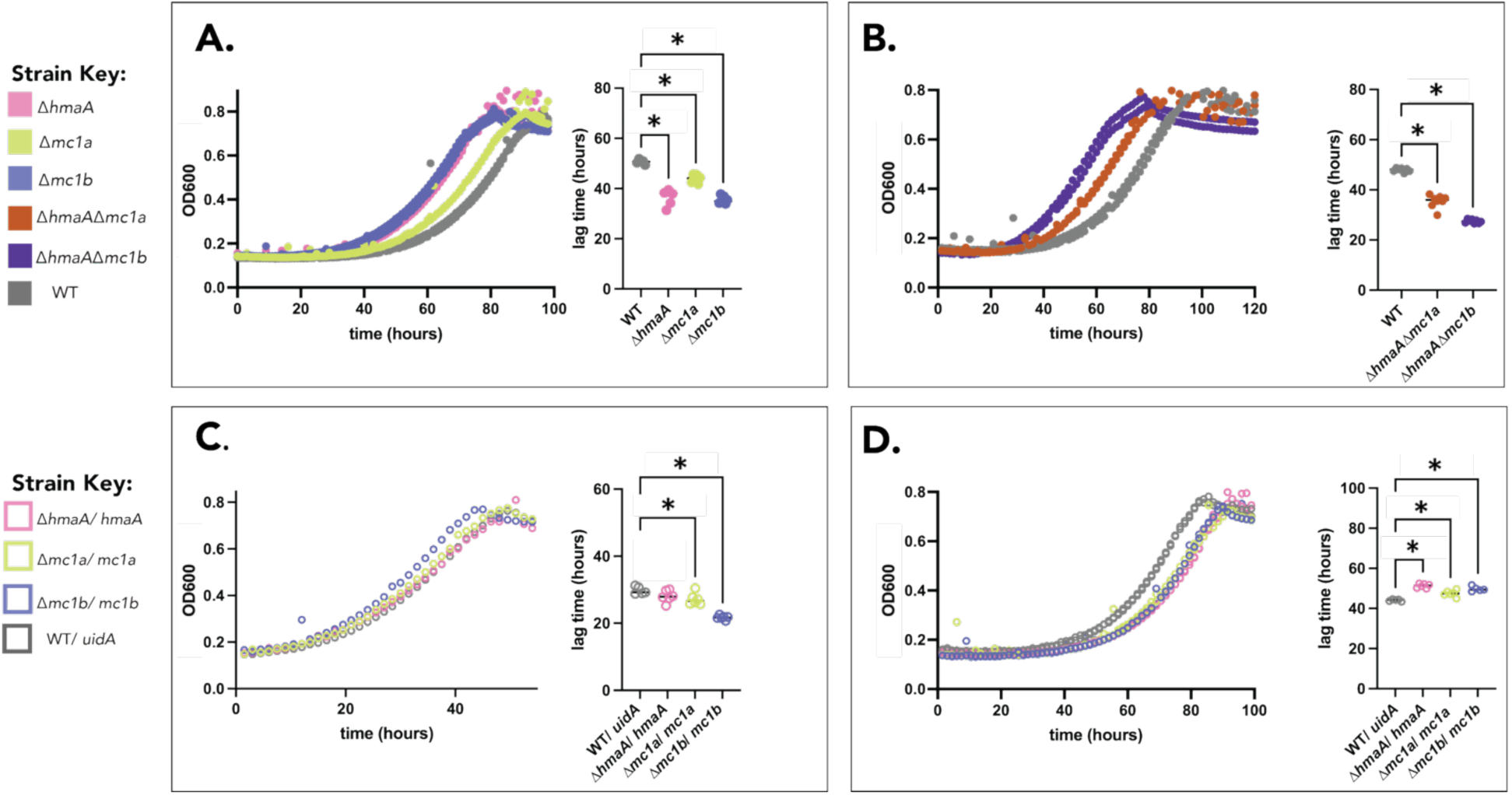
DNA-binding proteins (DBPs) affect timing of exit from extended stationary phase in *Methanosarcina acetivorans*. **A.** Growth curves of the single DBP deletion mutants (colored closed circles) in comparison to the parental strain (WWM60, denoted as wildtype or WT in gray closed circles) measured as a change in optical density (at 600 nm; OD600). Strains were incubated for 21 days incubation in stationary phase and inoculated in high-salt (HS) minimal medium supplemented with 50mM trimethylamine as the growth substrate. **B.** Growth of double DBP deletion strains (colored circles) compared to the WT strain (gray circles) under the same conditions as panel **A**. **C.** Growth of strains overexpressing DBPs in the respective DBP-deletion background (colored open circles) compared to the parental strain expressing an equivalent amount of *uidA* (encoding β-glucuronidase) (gray open circles). Growth curves in **C** are conducted after incubation of strains for 1 day in stationary phase. **D.** Growth curves of the same strains in panel **C** after incubation for 21 days in stationary phase. Growth media was supplemented with 2 μg/mL puromycin and 100 μg/mL tetracycline to select for maintenance of the expression vector and to fully induce expression of the gene, respectively (see Methods). Two representative growth curves for each strain are shown. The lag time (in hours) was quantified for least three replicates of each strain shown. The horizontal line represents the mean of the replicates. Statistically significant differences relative to the WT control strains, (p<0.05) based on a one-way ANOVA with Dunnett’s multiple comparisons test, are depicted as follows: * denotes p ≤ 0.05. See Table 3 for quantification of the growth parameters shown in this figure.

**Figure 5:**
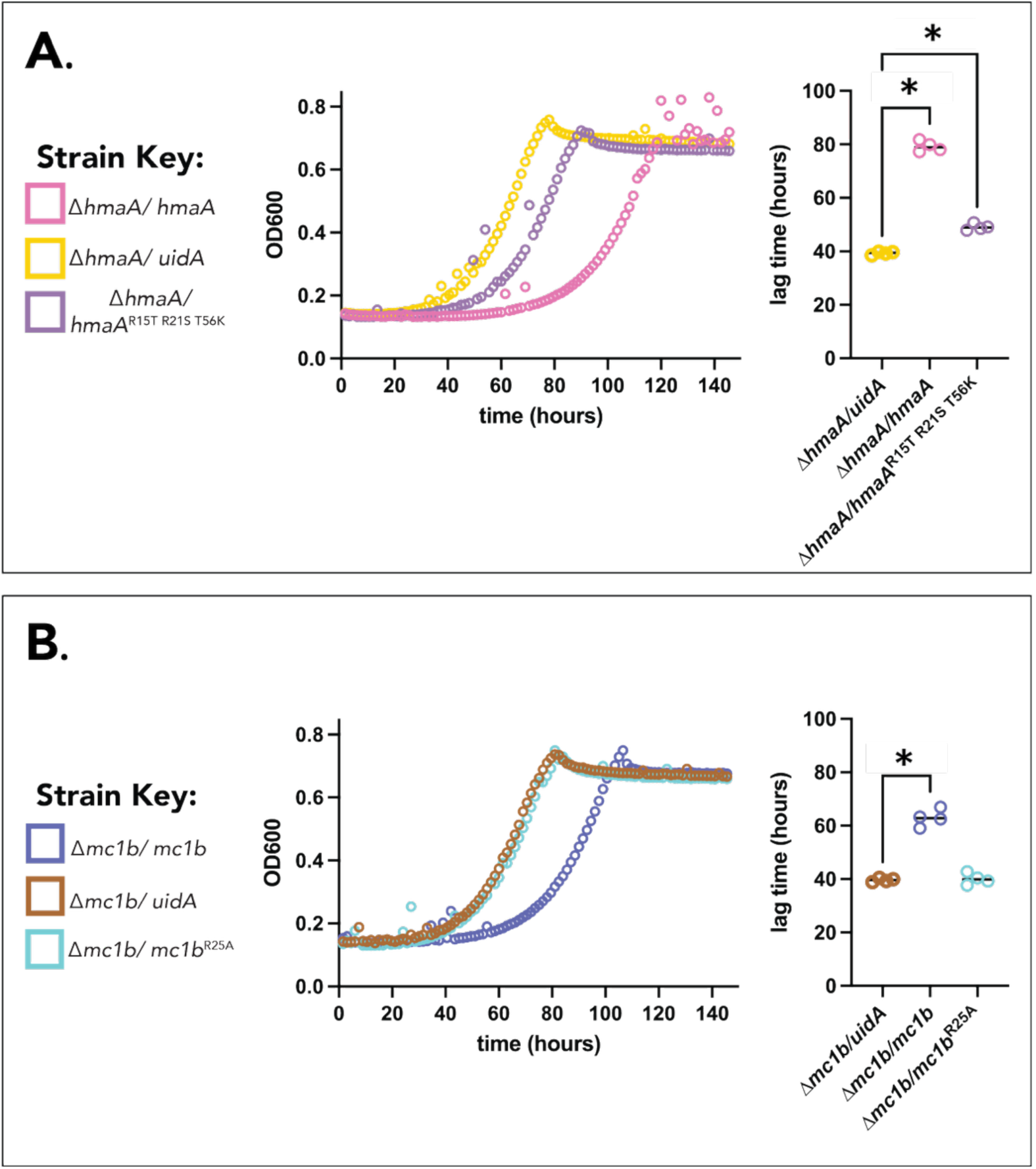
Abrogating DNA-binding residues in DBPs shortens exit from extended stationary phase in *Methanosarcina acetivorans*. **A.** Growth curves of *ΔhmaA-*derived strains expressing either HmaA:FLAG (pink open circles), HmaA ^R15T^ ^R21S^ ^T56K^: FLAG (lavender open circles) or UidA (yellow open circles) following 28 days incubation in stationary phase. **B.** Growth curves of *Δmc1b-*derived strains expressing either Mc1b:FLAG (blue open circles), MC1b ^R25A^: FLAG (brown open circles) or UidA (cyan open circles) following 28 days incubation in stationary phase. For both panels, a representative growth curve for each strain is shown. Growth media was supplemented with 2 μg/mL puromycin and 100 μg/mL tetracycline to select for maintenance of the expression vector and to fully induce expression, respectively (see Methods). The lag time (in hours) was quantified for least three replicates of each strain shown. The horizontal line represents the mean of the replicates. Statistically significant differences relative to the WT control strains, (p<0.05) based on a one-way ANOVA with Tukey’s multiple comparisons test, are depicted as follows: * denotes p ≤ 0.05. See Table 4 for quantification of the growth parameters shown in this figure.

**Table 3:** Growth data for stationary exit of DBP deletion and over-expression strains shown in Figure 4. Column 2 denotes the length of incubation in stationary phase for each strain before strains were inoculated into fresh media (1:10 dilution) and re-growth was tracked. Doubling time and lag time are reported for each strain upon re-growth in new media. Data represent at least three replicates. Mean values and standard deviations for each growth parameter are denoted. For values that are significantly different from the WT strain for a given condition (one-way ANOVA, Dunnett’s multiple comparisons test), percentage change from the WT is reported; sign (+ or -) indicates whether the mutant showed an increase or reduction relative to WT, respectively. All growth curves were conducted in a 24-well plate with an automated plate reader (see Methods) using high-salt (HS) minimal medium supplemented with 50mM TMA. For strains bearing an over-expression plasmid for complementation of a given DBP *in trans*, the media was supplemented with 2μg/mL puromycin and 100 μg/mL tetracycline, for maintenance of the plasmid and full induction of expression, respectively.

| strain | experiment conditions | doubling time (hours) | significantly different than WT (p<0.05)? | lag time (hours) | significantly different than WT (p<0.05)? |
| --- | --- | --- | --- | --- | --- |
| WT | 21 days stationary | 17.6±0.4 | n/a | 50.6±0.9 | n/a |
| $\Delta hmaA$ | | 18.2±1.3 | no | 36.6±3.2 | yes, -27.6% |
| $\Delta mc1a$ | | 20.3±2.7 | yes, +15.3% | 43.9±1.6 | yes, -13.3% |
| $\Delta mc1b$ | | 18.0±0.3 | no | 35.5±1.6 | yes, -30.0% |
| WT | 21 days stationary | 19.8±0.6 | n/a | 47.9±0.8 | n/a |
| $\Delta hmaA\Delta mc1a$ | | 20.2±1.4 | no | 35.4±2.6 | yes, -26.1% |
| $\Delta hmaA\Delta mc1b$ | | 17.7±0.8 | yes, -10.3% | 27.4±0.8 | yes, -42.7% |
| WT/ *uidA* | 1 day stationary | 16.3±0.7 | n/a | 29.8±1.2 | n/a |
| $\Delta hmaA/ hmaA$ | | 18.0±1.5 | no | 28.1±1.8 | no |
| $\Delta mc1a/ mc1a$ | | 14.7±1.5 | no | 27.2±1.8 | yes, -8.8% |
| $\Delta mc1b/ mc1b$ | | 15.5±0.5 | no | 21.7±0.6 | yes, -27.1%% |
| WT/ *uidA* | 21 days stationary | 17.74±1.5 | n/a | 44.1±0.5 | n/a |
| $\Delta hmaA/ hmaA$ | | 18.2±0.3 | no | 51.3±1.1 | yes, +16.3% |
| $\Delta mc1a/ mc1a$ | | 18.4±1.3 | no | 47.5±1.5 | yes, +7.7% |
| $\Delta mc1b/ mc1b$ | | 16.9±0.4 | no | 49.7±1.2 | yes, +12.8% |

**Table 4:** Growth data for stationary exit of DBP complementation strains shown in Figure 5. Column 2 denotes the length of incubation in stationary phase for each strain before strains were inoculated into fresh media (1:10 dilution) and re-growth was tracked. Doubling time and lag time are reported for each strain upon re-growth in new media. Data represent at least three replicates. Mean values and standard deviations for each growth parameter are denoted. For values that are significantly different from the WT strain for a given condition (one-way ANOVA, Tukey’s multiple comparisons test), percentage change from the WT is reported; sign (+ or -) indicates whether the mutant showed an increase or reduction relative to WT, respectively. All growth curves were conducted in a 24-well plate with an automated plate reader (see Methods) using high-salt (HS) minimal medium supplemented with 50mM TMA. The media was supplemented with 2μg/mL puromycin and 100 μg/mL tetracycline, for maintenance of the complementation plasmids and full induction of expression, respectively.

| strain | experiment conditions | doubling time (hours) | significantly different than *uidA* control? (p<0.05)? | lag time (hours) | significantly different than *uidA* control? (p<0.05)? |
| --- | --- | --- | --- | --- | --- |
| *ΔhmaA/uidA* | exit after 1 day stationary, standard conditions | 13.7±0.5 | n/a | 13.4±0.4 | n/a |
| *ΔhmaA/hmaA* |  | 15.9±0.8 | no | 14.0±0.7 | no |
| *ΔhmaA/hmaA*<sup>R15T R21S T56K</sup> |  | 13.9±0.9 | no | 13.8±2.0 | no |
| *Δmc1b/uidA* | exit after 1 day stationary, standard conditions | 14.3±2.1 | n/a | 14.0±1.2 | n/a |
| *Δmc1b/mc1b* |  | 15.1±0.2 | no | 12.8±0.2 | no |
| *Δmc1b/mc1b*<sup>R25A</sup> |  | 13.3±0.6 | no | 13.9±0.4 | no |
| *ΔhmaA/uidA* | exit after 28 days stationary, standard conditions | 15.8±0.3 | n/a | 39.3±0.6 | n/a |
| *ΔhmaA/hmaA* |  | 19.6±0.8 | yes, +24.1% | 79.3±2.0 | yes, +101.5% |
| *ΔhmaA/hmaA*<sup>R15T R21S T56K</sup> |  | 17.5±1.0 | no | 49.1±1.2 | yes, +24.8% |
| *Δmc1b/uidA* | exit after 28 days stationary, standard conditions | 16.7±0.6 | n/a | 39.7±0.7 | n/a |
| *Δmc1b/mc1b* |  | 19.6±2.1 | yes, +17.7% | 62.9±3.2 | yes, +59.3% |
| *Δmc1b/mc1b*<sup>R25A</sup> |  | 17.8±1.4 | no | 40.1±2.1 | no |

## Discussion

Even though histones and NAPs are primarily used for genome compaction in eukaryotes and bacteria, respectively, they also have additional roles in the cell. Representatives of these classes of proteins often co-occur in archaeal genomes, and it is unclear whether these proteins jointly compact the genome or perform alternate functions. Our genetic analyses show that the histone HmaA in *M. acetivorans* is not only dispensable but does not seem to contribute to fitness under routine laboratory conditions (Fig.1 and 3, Fig. S9). These observations deviate from a previous study conducted in *M. mazei*, perhaps because of differences in the mutant construction strategy used. Here, we generated a clean knockout using our Cas9-based genome editing technique; meanwhile, the corresponding mutant in *M. mazei* has a *pac* cassette under the control of the strong *mcr* promoter (P*mcrB*-*pac*) in place of the HMm locus^18^. It is plausible that the cost of protein overexpression or the presence of puromycin in the growth medium contributed to the growth phenotype of the Δ*hmm::PmcrB-pac* mutant. Alternately, we cannot rule out the possibility that the cellular function of histones is variable across *Methanosarcina* species. We were also able to generate mutants lacking one of the two copies of MC1 and delete *hmaA* in each of these backgrounds. However, despite multiple attempts, we could not obtain a mutant lacking both copies of *mc1* (Table S2). These data are consistent with an essential role for the MC1 proteins, perhaps in genome compaction. However, the global transcriptomic profiles (Fig.2, Table S3) and growth phenotypes (Fig. 3 and 4) suggest that the cellular function of the two MC1 proteins are not completely overlapping, which indicates that they have undergone partial neofunctionalization post-duplication. Since MC1a is more highly expressed and lacks signatures of sequence-specific DNA-binding, it is likely the primary driver of genome compaction in *M. acetivorans.* While MC1b might take on the role of genome compaction in the absence of MC1a, our data suggests that its primary function is likely more akin to a transcription factor. While the MC1b regulon is not enriched in any single functional category of genes, it is plausible that MC1b is a master regulator of many different regulons as 16 genes predicted to bind DNA and/or regulate transcription are differentially expressed in the Δ*mc1b* mutant (Table S7). At least one copy of *mc1* is conserved throughout the family *Methanosarcinaceae*, however several strains, including *M. acetivorans,* encode multiple copies (Fig. S15). Thus, we posit that duplication led to the emergence of unique functions in each paralog. Notably, Čuboňová and colleagues hypothesized that a similar dynamic has arisen between the two *Thermococcus kodakarensis* histone paralogs, which are individually dispensable (albeit, with a fitness cost) but cannot be deleted in tandem^34^.

Intriguingly, many of the DBP mutants in *M. acetivorans* seemed to have a growth advantage. These observations are in contrast to previous reports on DBPs in archaea. In halophilic archaea, deletion of histones leads to reduced fitness, even under optimal growth conditions^16^. Similarly, in *Solfolobus islandicus,* knockdown of the Alba family protein Sis10b resulted in slow growth compared to the parental strain^35^ In *Thermococcus kodakarensis,* deletion of the histone paralog, HtkA, abolishes transformation competency whereas a HtkB knockout has slower growth and lower fitness^34^. However, it has been routinely shown that deletion of NAPs in bacteria can have a growth advantage. For example, deletion of *hns* in *Escherichia coli* leads to a growth advantage at low pH and high osmolarity^36^ and a *hns* point mutant, *hns-66,* confers a growth advantage after prolonged stationary phase^37^. Similarly, *E. coli* strains lacking *fis* have increased growth rate and biomass yield on acetate minimal media^38^. Thus, it stands to reason that the fitness impacts of prokaryotic DBPs are specific to the individual organism and the growth conditions and cannot be broadly generalized.

Our finding that DBPs have growth-phase specific phenotypes are also consistent with prior art. For example, deletion of histone *hpyA* in *Halobacterium salinarum* results in a growth-phase dependent morphological phenotype^39^. In *S. islandicus,* Lrs14 has a growth-phase and stress-specific expression pattern^40^ and, in *H. salinarum,* DBPs levels increase during stationary phase. Altogether, it seems likely that DBPs in archaea might play a role in protecting genomic DNA during periods of sub-optimal conditions. While the precise molecular mechanisms of DBP function under these conditions remain to be elucidated, we suggest that DBPs likely benefit the organism in promoting long-term survival, perhaps by delaying growth until the organism encounters an environment that is more consistently favorable for growth. Indeed, as most microbes in their natural environment are not experiencing extended periods of exponential growth, this may explain why DBPs such as histone and MC1 are highly conserved across the archaeal tree^8^, yet, in some organisms, easily dispensable under laboratory growth conditions.

Most studies on archaeal histones have focused on their role in chromatinization of the genome to highlight the ways in which these proteins mirror their eukaryotic counterparts. From this perspective, the dispensability of histones in archaea is peculiar and confounding. Even though this eukaryocentric view is evolutionarily sound, it disregards the cytological features of extant archaea, which are a lot like bacteria. Archaea lack nuclei and have a single circular chromosome that is likely compacted into a nucleoid-like structure. Like bacteria, that often encode multiple NAPs at once, archaea too encode histones and suite of NAPs. From this standpoint, the dispensability of histones (or any other NAP) in some archaea is not at all unprecedented. Many NAPs in bacteria, like H-NS and DPS, are not essential either^41,42^. Thus, we propose that histones in archaea should be considered as NAPs, with context-dependent functions that apply based on the lifestyle of the organism and other NAPs present within that system. We also posit that histones have no unified function across the archaea. In some systems they may be a core component of chromatin architecture, as has been previously observed^13,14^, while in other systems they may function more like transcriptional regulators^16^, play a role in stress response^18^, or mediate quiescence. In this way, archaeal histones join the ranks of most other prokaryotic NAPs, which often vary widely in function across the tree of life.

## Materials and Methods

### Structure alignment of DNA-binding proteins

The *Methanothermus fervidus* HmfB crystal structure (PDB: 1a7w) or *Methanosarcina thermophila CHT155* MC1 solution structure (PDB: 2khl, model 1.1) were aligned with the AlphaFold2^43^ predicted structure for *Methanosarcina acetivorans* HmaA (AF-Q8TSX3-F1_model_v4.pdb, AlphaFold Protein Structure Database) and *M. acetivorans* MC1a (AF-Q8TK06-F1-model_v4.pdb, AlphaFold Protein Structure Database) or *M. acetivorans* MC1b (AF-Q8THJ8-F1_model_v4.pdb, Alpha Fold Protein Structure Database), respectively, using the matchmaker command in UCSF ChimeraX^44^ (version 1.7.1). The Root Mean Square Displacement (RMSD) output is reported based on the pruned structure. For the HmfB-HmaA alignment, the pruned structure represents 66 of 67 possible atoms. For the MC1-MC1a alignment, the pruned structure represents 71 of 93 possible atoms. For the MC1-MC1b alignment, the pruned structure represents 72 of 93 possible atoms.

### Media and culture conditions

*Methanosarcina acetivorans* strains were cultivated at 37 °C without shaking in bicarbonate-buffered high salt (HS) medium^45^ supplemented with either 50 mM trimethylamine hydrocholoride (HS-TMA), 125 mM methanol (HS-MeOH), or 40mM sodium acetate (HS-acetate) as growth substrates. Prior to sterilization, growth substrates were added along with NH_4_Cl (final concentration, 19 mM), Cysteine-HCl (final concentration, 2.8 mM), Na_2_S.9H_2_O (final concentration, 0.4 mM), vitamin mix, and trace elements solutions, as described previously^46^. Cultures were grown in sealed Balch tubes with N_2_/CO_2_ (80:20) in the headspace at 8-10 psi. *Escherichia coli* strains were cultivated at 37 °C shaking at 250 RPM in lysogeny broth (LB) supplemented with 20 µg/mL chloramphenicol and/or 25 µg/mL kanamycin.

### Plasmid construction

Plasmids for CRISPR-Cas9-mediated genome editing were cloned as previously described^26^. Single guide (sg)RNAs were designed using the CRISPR site tool in Geneious Prime (version 2024.0.5, https://www.geneious.com); sequences are provided in **Table S10**. PCR fragments containing the sgRNA scaffold were assembled into pDN201 linearized with *AscI* as previously described^26^. The appropriate homologous repair (HR) template was assembled into the sgRNA containing derivative of pDN201 at the *PmeI* site. For complementation plasmids, PCR fragments containing genes of interested were amplified from the *M. acetivorans* genome or from commercially synthesized gene fragments (gblocks) containing the desired point mutations, as applicable. When applicable, epitope tags were added to gene fragments using a primer containing the desired tag sequence. Primer information can be found in **Table S10**. PCR fragments containing genes of interest, and tags, when appropriate, were fused to the appropriate tetracycline-inducible promoter by linearizing either pJK027a or pJK029a with *NdeI* and *HindIII.* Plasmids were verified by Sanger sequencing. For expression in *M. acetivorans,* pJK027A/ pJK029A derivative plasmids were co-integrated with pAMG40. Co-integration was conducted using the Gateway BP Clonase II Enzyme Mix and verified by a diagnostic restriction endonuclease digest. All cloning was performed in WWM4489, an *E. coli* DH10B derivative, which was transformed via electroporation at 1.8 kV using an *E. coli* Gene Pulser. *E. coli* strains carrying the cointegrated vector were cultured with 10 mM Rhamnose to induce copy number as previously described^47^. All primers and plasmids used in this work are cataloged in **Tables S10** and **S11**, respectively.

### Generation of *M. acetivorans* deletion mutants

Transformation of *M. acetivorans* was conducted as described previously^48^. In brief, 20 mL cultures of *M. acetivorans* were grown in HS media supplemented with 50mM TMA (HS-TMA) as the growth substrate. Transformants were plated on solidified HS-TMA + 1.5% agar medium supplemented with 2 µg/mL puromycin as a selection agent. Plates were incubated at 37 °C in an anaerobic incubator with H_2_S/CO_2_/N_2_ (1000 ppm:20:balance) in the headspace. For complementation strains, colonies were screened by PCR and sequenced by Sanger sequencing to verify presence of the desired plasmid construct. For deletion strains, individual colonies were picked into liquid medium, grown to saturation, and then screened via PCR at the chromosomal locus for the desired mutation. Screening primers are listed in **Table S10**. Colonies that were identified as having the desired deletion were then plated on solidified HS-TMA + 1.5% agar medium supplemented with 20 µg/mL 8ADP to cure the plasmid. Single colonies were then picked and screened via PCR to confirm absence of the *pac* (puromycin resistance) gene on the mutagenic plasmid (see **Table S10** for primers). All deletion mutants in this study were re-sequenced via whole-genome Illumina sequencing (see below) to ascertain the presence of the desired chromosomal deletion, as well as to screen for any possible suppressor mutations. A list of strains used in this study is provided in **Table S12**. In the case that a desired DBP mutant was unable to be recovered (e.g. *Δmc1aΔmc1b*), the transformation of the editing plasmid was repeated at least two more times and colonies were screened as described above. In parallel, and within the same background where deletion of the DBP was attempted, an editing plasmid targeting a neutral locus (MA2520, orotate phosphoribosyltransferase-like protein), was also transformed. Colonies were then screened for deletion of the neutral locus. A mutant was considered unable to be generated when deletion of the neutral locus, but not the desired DBP locus, was observed in multiple biological replicates. A summary of these experiments can be found in **Table S2**.

### DNA extraction and whole genome re-sequencing

10mL cultures were grown to saturation and DNA was extracted and processed as previously described^46^. Genomic DNA was sequenced at the SeqCenter (Pittsburgh, PA) via Illumina paired-end reads. The resulting sequencing reads were subsequently mapped to the *M. acetivorans* reference genome using Breseq^49^ version 0.35.5 (default parameters). In strains cured of the editing plasmid, *M. acetivorans* C2A was used as the reference genome. In strains that were re-sequenced while bearing the editing plasmid, *M. acetivorans* strain WWM60 concatenated with the applicable editing plasmid (e.g. pAB13) was used as the reference. All mutations identified via this analysis are reported in **Table S1.**

### Cultivation for growth assays in Balch tubes

Growth curves in Balch tubes, as delineated in **Table 1**, were performed by pre-culturing strains in 10 mL HS media at 37 °C without shaking in sealed Balch tubes with N_2_/CO_2_ (80/20) in the headspace at 8-10 psi. Media was supplemented with either 50 mM TMA, 125 mM MeOH, or 40 mM acetate, as indicated in **Table 1**. For growth experiments, Balch tubes containing sterile media were inoculated with 0.5 mL of the pre-culture at late exponential phase (OD600 ∼0.6-0.9). In experiments with multiple strains, inocula were normalized to account for any OD600 differences between strains. Growth of four independent biological replicates were monitored for each strain by monitoring the change over time in optical density of cultures at 600 nm using a UV-Vis Spectrophotometer (Genesys 50, Thermo Fisher Scientific, Waltham, MA, USA). Readings were “blanked” with an uninoculated, sterile tube containing 10 mL of the corresponding medium. Growth rates were determined by finding the slope of the best linear fit of the log-transformed OD600 versus time plot. The ‘best linear fit’ was determined by selecting the linear model with the maximal R^2^ value (minimum cut-off ≥ 0.98) with a minimum four points. Doubling times were calculated from growth rates by dividing the natural log of 2 by the growth rate. Lag time was calculated by solving the best linear fit model for time x when y is set equal to the log-transformed initial OD600 observed at the beginning of the experiment.

### Cultivation for high-throughput microplate reader growth assays

Growth curves conducted in a high-throughput microplate reader format were performed by using a modified HS-TMA medium where sodium bicarbonate was removed from the medium and the medium was instead buffered by 50 mM 1,4-piperazinediethanesulfonic acid (PIPES) adjusted to a final pH of 6.8 (HS-TMA-PIPES). All other components of the medium remained the same as described above. Strains were pre-cultured until they reached stationary phase in 10 mL HS-TMA-PIPES in hermetically sealed Balch tubes with pure N_2_ in the headspace at 8-10 psi. Within 24 hours of reaching stationary phase (unless otherwise specified), 1 mL of precultures were then inoculated into Balch tubes containing fresh HS-TMA-PIPES media; for growth curves with multiple strains, inocula were normalized to account for any differences in the OD600 between strains. Freshly inoculated Balch tubes were then brought inside of an anaerobic chamber (Coy Laboratory Products, Grass Lake, MI, USA) and decapped. The contents of each Balch tube were then mixed by gently pipetting up and down with a 10 mL pipette at least 10 times to ensure homogeneous distribution of the cultured cells within the liquid medium. 1.8 mL of the inoculated cultures were then dispensed into each well of a sterile, anaerobic 24-well flat-bottomed tissue culture plate (cat. #353047, Corning Life Sciences, Durham, NC, USA). For growth curves with multiple strains, strains were assigned well locations via random number generator, so as to mitigate any edge-effects or global position bias in the growth of an individual strain. In any given growth curve, at least four replicate wells were used for a given strain. Plates were tightly sealed with optical film (cat #. 2921-7800, USA Scientific, Ocala, FL, USA) and then incubated in a microplate reader (Agilent BioTek Epoch 2, Agilent, Santa Clara, CA, USA) at 37°C. The microplate reader was programmed as follows: experiment type-24-well kinetic assay, kinetic activity-shake before each read at 425cpm (3mm, speed: fast), primary wavelength-600nm. For growth curves where readings were taken every hour, the kinetic interval was set to 1 hour and the shake interval was set to 59 minutes. For growth curves where readings were taken every 1.5 hours, the kinetic interval was set to 1 hours, 30 minutes and the shake interval was set to 1 hr, 29 minutes. Growth data were analyzed with a custom MATLAB script which determined a best-fit linear model from natural log-transformed growth data. The ‘best linear fit’ was determined by selecting the linear model with the maximal R^2^ value (minimum cut-off ≥ 0.99) with a minimum eight points. Doubling time and lag time were then calculated as described above. Upon conclusion of the growth curve, if defects were observed in the seal integrity around a given well, data from that well was not considered in analysis. For growth experiments with strains bearing a complementation plasmid *in trans*, the growth media were supplemented with 2 µg/mL puromycin for plasmid maintenance and 100 µg/mL tetracycline for full induction of the inducible promoter.

### Electron microscopy and cell morphology analysis

Strains were precultured in HS-TMA. At late exponential phase, cells were fixed in 4% paraformaldehyde for 25 minutes, then washed once in 3x phosphate-buffered saline (PBS). Cells were then settled onto plasma-treated coverslips, post-fixed with osmium tetraoxide for 1 hour, then ethanol dehydrated for 30 minutes. Samples were then placed in the critical point dryer for at least two hours, sputter coated with platinum for 30 minutes, and imaged on the scanning electron microscope (Phenom Pharos, Nanoscience Instruments Inc.). Cell width was measured in FIJI^50^ by drawing a horizontal line from edge to edge of each cell and using the “measure” function to determine width in microns. 100 individual cells were measured for each strain. Cells were excluded from measurement if they were clumped with other cells or displayed visible extrudate suggesting that membrane integrity had been compromised.

### RNA extraction, sequencing, and analysis

10 mL cultures of the parental strain (‘WWM60’) and DBP single knockout strains *(ΔhmaA, Δmc1a, Δmc1b*) were grown in quadruplicate on HS-TMA at 37 °C. When cultures reached mid-exponential phase (OD600 0.5-0.6), 1mL culture was removed and processed as described previously^46^. Samples were sent to SeqCenter (Pittsburgh, PA) for Illumina paired-end sequencing. Transcriptome analysis was performed using the KBase bioinformatics platform as previously described^46,51^. Briefly, raw reads were aligned to the *M. acetivorans* C2A reference genome using Bowtie2 (v.2.3.2); aligned reads were assembled using Cufflinks (v.2.2.1) and fold change, significance values, and a principal component analysis were calculated using DESeq2 (v.1.20.0). Default parameters were used for all programs. We designated a gene as differentially expressed (DE) from the WT strain if it exceeded a log_2_ fold change of ±0.1 and a q-value of ≤0.05. DESeq2 data for all mutant strains, with respect to the WT strain, is listed in **Table S3.** Raw transcript reads have been deposited in the Sequencing Read Archive (SRA), project number PRJNA1365308.

Genes designated as DE in each mutant strain were analyzed for COGs (clusters of orthologous genes^52,53^) using COGclassifier^54^ with default parameters. Promoters regions of DE genes were assessed for enrichment of DNA sequence motifs using the MEMEsuite platform^55^. Sequences of promoter regions were obtained by collecting 500 base pairs of upstream DNA sequence from each gene designated as DE. These sequences were compiled into a FASTA file which was run through the MEME motif discovery program in differential enrichment, any number of repetitions (-anr), and a motif cap of 5. The other parameters were default. The control sequences for differential enrichment mode was a set of 500 sequences of length 500bp subsampled at random from the *M. acetivorans* C2A genome using a custom script. The MEME suite program was run locally on a laboratory computer via Docker Desktop.

### Growth assays with stressors

For assessment of growth with stressors, as depicted in **Figure 3** and **Table 2**, strains were cultivated as described for high-throughput microplate reader growth assays, with the following exceptions: for growth with mitomycin C, growth media were supplemented with 1 µg/mL mitomycin C (Sigma Aldritch, cat# M4287). For growth at high temperature, strains were pre-acclimated by culturing in HS-TMA-PIPES, in Balch tubes submerged in a 40 °C water bath, for three passages prior to inoculation and growth in the microplate reader at 40 °C. For UV shock, strains were pre-cultured in HS-TMA-PIPES in Balch tubes as described above, then exposed to UV light in alignment with methods previously described^18^. Briefly, exponentially growing strains (OD600 = 0.6-0.9) were brought into the anaerobic chamber and decapped, and the 10 mL pre-culture volume was dispensed into open, anaerobic petri dishes. The suspended cultures were gently stirred, then subjected to 253 nm UV light from a handheld UV lamp (iClevr, Newark, CA, USA) at 20 cm for 3 minutes. Following irradiation, cultures were gently stirred to ensure homogenous mixing of cells throughout the culture, then a 3 mL volume, normalized to account for any differences in OD between strains, was inoculated into 10 mL fresh HS-TMA-PIPES media. Recovery and subsequent growth were then monitored in 24-well plates as described above.

### Assessment of strain growth following extended stationary phase incubation

For experiments examining stationary phase exit, strains were pre-cultured in HS-TMA-PIPES in Balch tubes as described above. During pre-culturing, OD600 was monitored to ensure strains were synchronized in terms of growth phase. Once strains reached saturation and entered stationary phase, as defined by the point where OD600 stops monotonically increasing, strains were continuously incubated at 37 °C for the amount of time specified in **Figs. 4 and 5** and **Tables 3 and 4**. After incubation in stationary phase, 1 mL of each pre-cultured strain, normalized to account for any differences in OD600 between strains, was inoculated into fresh HS-TMA-PIPES medium and subsequent recovery and growth was monitored via microplate reader growth curves as described above. For strains bearing complementation plasmids, both the pre-culture media and media for the growth curve experiment was supplemented with 2 µg/mL puromycin for plasmid maintenance and 100 µg/mL tetracycline for full induction of the inducible promoter. During pre-culture, Balch tubes containing media with tetracycline were wrapped in foil for protection from ambient light.

### Microscopy assay for intact cell membranes

1 mL of culture was removed from a Balch tube and spun down at 10,000g in a bench-top microcentrifuge. 980 µL supernatant was removed and the cell pellet was resuspended in the remaining 20 µL media. 5 µL of this resuspension was mounted onto a glass slide for imaging. Imaging was conducted using a Zeiss AxioImager M2 microscope. Brightfield images were captured using differential interference contrast (DIC) and factor F_420_ autofluorescence was captured in the DAPI channel. At least three random fields comprising at least 250 cells were imaged for each strain under each condition. Cells visible in each channel were quantified in FIJI for each image as follows: image thresholds were adjusted to select cells and then converted to binary. The binary > fill holes and binary > watershed commands were run to ensure selection of individual cells. The analyze > analyze particles command was run with the size selection from 10 to infinity; otherwise, default parameters were used. The cell count was determined by the ‘count’ output. The percentage of intact cells was determined by dividing the count of cells in the DIC channel to the count of cells in the DAPI channel for each field of cells.

### Immunoblotting of FLAG-tagged proteins

Strains bearing over-expression plasmids for FLAG-tagged proteins were grown in 10mL HS-TMA medium grown to saturation. 2 µg/mL puromycin was added to maintain the plasmid. When appropriate, 100 µg/mL tetracycline was added for full induction of gene expression. Cell lysates were prepared and processed as described previously^46,51^. Total protein concentration in cell lysates were quantified using Bradford reagent (Sigma-Aldrich, St Louis, MO) with bovine serum albumin (BSA) standards, per the manufacturer’s instructions. Volumes of cell lysate containing approximately equal quantities of protein were mixed with 2x lammeli loading buffer (β-mercaptoethanol added), then loaded onto a 4-20% gradient mini-Protean TGX gel (Bio-Rad). In gels with dilution series (**Fig S13C-E**), cell lysates were normalized to a concentration of 10 mg/mL protein (HmaA and HmaA^R15T^ ^R21S^ ^T56K^ samples) or 5 mg/mL protein (MC1b and MC1b^R25A^ samples). These were then diluted in series as indicated in the lane key of **Fig S13** and equal volumes were loaded on the gel as described above. Following separation on the gel, proteins were transferred to a Trans-Blot Turbo Mini 0.2 µM PVDF membrane using the Bio-Rad Trans-Blot Turbo Transfer System. FLAG-tagged proteins were visualized using monoclonal anti-FLAG M2-Peroxidase (HRP) antibody (Sigma-Aldrich, St Louis, MO) at a 1/30,000x dilution. Signal from the antibody was detected using Immobilon Western Chemiluminescent HRP Substrate (Millipore, Burlington, MA) with images collected via a ChemiDoc MP Imaging System (Bio-Rad, Hercules, CA). Exposure time for each blot was 1 or 3 seconds, as indicated in **Fig S13**. For panel **E** of **Fig S13**, a clean paper towel was placed over the right half of the blot to mask the bright bands present on that side of the blot and allow for better visualization of the dimmer bands on the left side of the blot. The same blot, with no mask, is shown in **Fig S13D** with an exposure time of one second. As a loading control for each Western blot, duplicate samples were loaded onto a second gel of the same composition and layout; this gel was stained and protein bands were visualized using GelCode Blue (ThermoFisher, cat #24590). Loading controls are depicted in **Fig S13A** and **S13C**.

### Bioinformatic screen for histone and *mc1* genes within the *Methanosarcinaceae*

Archaeal genomes from GTDB r.214.0 were downloaded, annotated, and checked for genome completeness as previously described^46,56^. Profile HMMs for histone (accession: NF043032.1) and MC1 (accession: PF05854.hmm) were obtained from NCBI and InterPro, respectively. The HMM Gene Search Tools were used with default parameters. All protein hits were counted and the number of hits was recorded for each genome. This information was overlaid on a tree of Methanosarcinaceae from the GTDB ar53_r214.tree using iTOL^57^.

## Data and code availability

Sequence reads for whole genome sequencing and RNA sequencing in this study have been uploaded to the Sequencing Reads Archive (SRA); project number PRJNA1365308. Custom scripts used for analysis in this manuscript can be found at https://github.com/alienorbask/growthcurve.

## Acknowledgements

We would like to thank all members of the Nayak Lab for valuable discussion, input, and support. We would like to thank Dr. Katie Shalvarjian for technical support in running the HMM search program and processing RNA-sequencing reads, Dr. Grayson Chadwick and Madison Williams for support with microscopy. Additionally, we thank Dr. Gabriella Pinter and Dr. Istvan Laukó with technical assistance in setting up the automated growth curve analysis pipeline. We also thank Dr. Dyche Mullins, Dr. Sam Lord, and Dr. Arthur Charles-Orszag for valuable discussion, input, and technical support with microscopy during the Archaeal Cell Biology Workshop, 2023. DDN acknowledges funding from the Searle Scholars Program sponsored by the Kinship Foundation, the Rose Hills Innovator Grant, the Beckman Young Investigator Award sponsored by the Arnold and Mabel Beckman Foundation, the Alfred P. Sloan Research Fellowship sponsored by the Sloan Foundation, the Simons Foundation Early Career Investigator in Marine Microbial Ecology and Evolution Award, and the Packard Fellowship in Science and Engineering sponsored by the David and Lucille Packard Foundation. DDN is a Chan-Zuckerberg Biohub – San Francisco Investigator. ASB would like to acknowledge funding from the NIH ‘Genetic Dissection of Cells and Organisms’ training program (award #5T32GM132022-03). DDN and ASB acknowledge funding from the Gordon and Betty Moore Foundation through project number GBMF 11481. The funders had no role in the conceptualization and writing of this manuscript or the decision to submit the work for publication.

